# Normative Data for an Expanded Set of Stimuli for Testing High-Level Influences on Object Perception: OMEFA-II

**DOI:** 10.1101/807446

**Authors:** Colin S. Flowers, Kimberley D. Orsten-Hooge, Barnes G.L. Jannuzi, Mary A. Peterson

**Affiliations:** Department of Psychology, University of Arizona, Tucson, Arizona, United States of America; School of Behavioral and Brain Sciences, The University of Texas at Dallas, Dallas, Texas, United States of America; Neuroscience and Cognitive Science Program, University of Arizona, Tucson, Arizona, United States of America; Cognitive Science Program, University of Arizona, Tucson, Arizona, United States of America

## Abstract

We present normative data for *bipartite displays* used to investigate high-level contributions to object perception in general and to figure-ground perception in particular. In these vertically-elongated displays, two equal-area regions of different luminance abut a central, articulated, vertical border. In *Intact* displays, a portion of a mono-oriented well-known (“familiar”) object is sketched along one side of the border; henceforth the “*critical* side.” The other side is the “*complementary side*.” We measured inter-subject agreement among 32 participants regarding objects depicted on the critical and complementary sides of the borders of *Intact* displays and two other types of displays: upright and inverted *Part-Rearranged* displays. The parts on the critical side of the border are the same in upright *Intact* and *Part-Rearranged* displays but spatially rearranged into a new configuration in the latter. Inter-subject agreement is taken to index the extent to which a side activates traces of previously seen objects near the central border. We report normative data for 288 regions near the central borders of 144 displays (48/type) and a thorough description of the image features. This set of stimuli is larger than an older “Object Memory Effects on Figure Assignment” (OMEFA) set. This new OMEFA-II set of high-resolution displays is available online (https://osf.io/j9kz2/).

---

A fundamental aspect of object perception involves determining whether a border between two regions in the visual field is a bounding contour of an object on one side, whether the border is *assigned to* one side, or *owned by* one side but not the other. When the border assignment occurs, the region on the side to which the border is assigned is perceived as a *figure* (i.e., an object) shaped by the border, whereas the other side is perceived as a locally shapeless *ground* (i.e., a background; e.g., [1 – 4]). Border assignment is influenced by figural priors – object properties associated with figures rather than backgrounds, including enclosure, symmetry, surroundedness, size, convexity, top-bottom polarity, lower region, contrast, and familiar configuration (e.g., [4 – 10]; for reviews: [1, 3, 11]).

The aforementioned figural priors are image characteristics. Another figural prior – *familiar configuration –* depends upon past experience rather than image characteristics (e.g., [2, 12, 13]; for review: [14, 15]). Effects of familiar configuration on figure assignment were demonstrated using vertically elongated *bipartite* displays like those in Figure 1, each consisting of two equal-area regions (one black, one white) meeting at a central, articulated, vertical border. The displays were designed so that the central border sketched a portion of a common mono-oriented object (that has a typical upright orientation) on one, “critical,” side and not on the opposite, “complementary,” side (see Figure 1A). This nominal difference between the two sides of the displays was affirmed in pilot experiments that revealed high inter-subject agreement regarding the common object resembled by the critical side of the border and low inter-subject agreement regarding any common object depicted on the complementary side of the border (cf., [12, 13, 16]).

**Figure 1.**
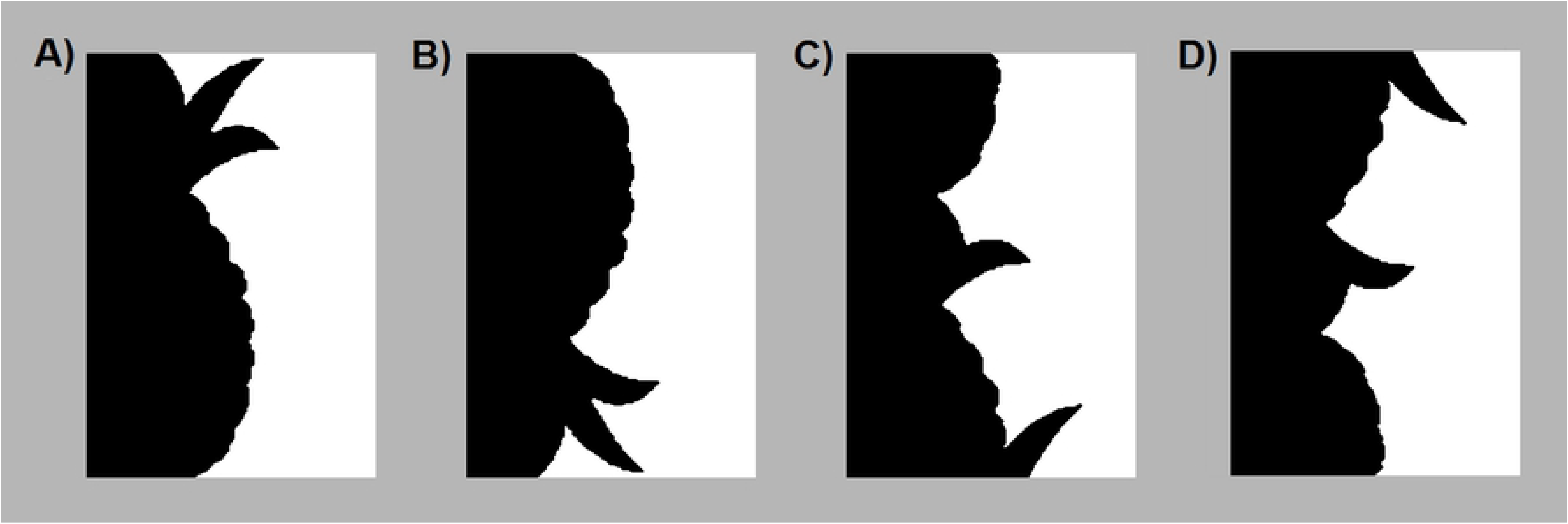
A sample bipartite stimulus in 4 configurations. In this figure, the critical side of the border is presented in black on the left of the central border. In the experiments the black/white contrast and left/right location of the critical side was balanced. A) Intact, B) Inverted, C) Upright Part-rearranged, D) Inverted Part-rearranged versions of the source stimulus, “Pineapple.” In experiments, the bipartite stimuli are presented on a medium gray backdrop so that the black and white sides contrast equally with the backdrop.

Peterson et al. [2, 12, 13, 16, 17] demonstrated effects of past experience by showing that the figure was more likely to be perceived on the critical side of the border in upright versus inverted versions of these displays (see Figures 1A & B). Image characteristics are held constant over a 180° orientation change but past experience is not because familiarity with mono-oriented objects is established by repeated exposure to them in their typical upright orientation; hence, inverted versions of mono-oriented objects are less familiar than upright versions. Peterson et al. [2, 16] demonstrated that these effects were due to the familiarity of configurations rather than of parts in experiments that showed that the figure was substantially more likely to be perceived on the critical side of the central border when the familiar configuration was sketched there in an intact form (i.e., its parts were arranged properly from top-to bottom; Figure 1A) than when its parts were spatially rearranged into a new, *Part-Rearranged*, configuration (Figure 1C). They reasoned that these effects manifested influences of object memories on figure assignment.

Previous experiments investigating effects of familiar configuration on figure assignment used ≤ 24 bipartite displays depicting a portion of an *Intact* familiar configuration on one side of the border with associated *Inverted Intact* and *Upright Part-Rearranged* versions. A set of stimuli originally used in experiments with brain-damaged participants, the “Object Memory Effects on Figure Assignment” (OMEFA) set has been used extensively [18 – 20]. Barense et al. [18, 21, 22] tested figure assignment with *Inverted Part-Rearranged* displays as well. Recently, we modified the borders of the OMEFA stimuli, producing high-resolution images; we also eliminated some items and added others. In this article, we report contemporary data on inter-subject agreement regarding the common objects resembled by the critical and the complementary sides of 144 bipartite displays in an expanded, fine-tuned, set of *Upright Intact* (N = 48), *Upright Part-Rearranged* (N = 48), and *Inverted Part-Rearranged* (N = 48 each) stimuli – the OMEFA-II stimulus set.

We used the Amazon Mechanical Turk (AMT) platform to gather contemporary norms regarding the familiar objects resembled by both sides of the border in the three types of displays. In what follows, we denote the stimuli by the name of the familiar configuration intended to be depicted by the *Upright Intact* displays (the “source” name) modified by display type. Individual participants viewed and responded to stimuli of all display types but, they saw a stimulus derived from a particular source stimulus in only one of the three display types. They viewed each stimulus for as long as they wished and listed up to three interpretations for each side of each bipartite display. *Inverted Intact* displays were not tested because when viewed for long periods of time, the inverted source object is easily recognized. However, we know that the critical side is assigned figure substantially and significantly less often in *Inverted Intact* displays then *Upright Intact* displays (e.g., [2]); therefore, the AMT norms for *Inverted Intact* displays would not be informative with regards to figure assignment processes.

Critical sides for which inter-subject agreement is high will be considered good depictions of portions of familiar objects. We expected to obtain high inter-subject agreement for the critical sides of many of the *Upright Intact* displays (that were designed to depict the source stimuli), but not their variants which were intended to control for image features while reducing or eliminating effects of familiar configuration. For objects with distinctive parts, we expected that the parts might support some degree of inter-subject agreement for the critical sides of *Part-Rearranged* displays, although not as much as for the critical sides of *Upright Intact* displays. We note that our method assesses explicit identification of familiar configurations, which we assume is related, but not identical, to implicit access to traces of previously seen objects that serves as a figural prior.

## Methods

### Participants

Potential participants had to meet the eligibility criteria of (a) having completed 1000 experiments or other data collection programs on AMT and (b) have achieved an approval rating of at least 95% (see [23]); 194 AMT participants met these criteria. Responses from 16 of these participants were excluded because they failed attention check trials (see Procedure); responses from four other participants were excluded because they were gibberish or non-words. Responses from the remaining 174 participants were analyzed.

Participants were compensated $1.50 to complete the task. Pilot tests showed that the tasks took no more than 10 minutes to complete (and could be completed much faster). Therefore, the estimated rate of pay was at the very least $9.00 per hour (above the US national minimum of $7.25 in 2015 and 2016 when these data were gathered).

### Stimuli

Bipartite displays are vertically elongated displays comprising two regions situated on the left and right sides of a central border. One region is black and the other white; they are presented on a medium gray background such that the black and white regions contrast equally with the background. Using AMT, we could not control exact luminance values on participants’ screen. We used pixel RGB values of: black = [0 0 0], white = [255 255 255], gray = [182 182 182]. These RGB values yielded luminance values of 0.12, 87.33, and 45.70 foot-lamberts respectively on the computers in our laboratory, though these surely differed for each individual AMT participant. The two regions are equated for area by equating the number of pixels in each region (mean % pixels on the critical side = 49.99% for Intact displays and 50.00% for Part-Rearranged displays; see Appendix A for image characteristics). We tested 48 bipartite displays with critical sides sketching Upright Intact versions of the 48 familiar *source configurations*, 48 upright Part-Rearranged versions of each of the source configurations, and 48 inverted Part-Rearranged versions of each of the source configurations. The 144 stimuli tested are listed in Table 1 and can be accessed online (https://osf.io/j9kz2/). Stimuli were 343 pixels high (H) and ranged from 111 to 350 pixels wide (W). AMT participants view the stimuli at different viewing distances and on screens with different sizes and different resolutions; hence, stimulus size was not matched across subjects in this experiment (although it was matched for the different display types individual participants viewed). However, the number of pixels in the stimuli uploaded to AMT was large enough that we could be reasonably confident that the stimuli were of sufficiently high resolution under these disparate conditions.

**Table 1.** Percent inter-subject agreement and difference scores for each side of three types of OMEFA-II bipartite stimuli: *Upright Intact, Upright Part-Rearranged*, and *Inverted Part-Rearranged*. The five columns under each type list (1-2) the interpretation with the highest inter-subject agreement for the critical side, (3-4) the interpretation with the highest inter-subject agreement for the complementary side, and (5) the critical – complementary difference. The first column denotes the source object, the object intended to be depicted on the critical side of the border of *Upright Intact* stimuli. Stimuli are ordered from top to bottom by percent inter-subject agreement regarding the interpretation for the critical side of the border. For the four objects listed in light grey at the bottom, either the interpretation with the highest inter-subject agreement was different from the source object or the critical – complementary difference was 0. The interpretations shown in bold for *Upright Part Rearranged* and *Inverted Part Rearranged* stimuli where neither side depicts an intact familiar object are interpretations that match the source object. Note that the Mickey Mouse stimulus is labelled as “Mickey” in the result and stimulus files.

| Source | Upright Intact |  |  |  |  | Upright Part-Rearranged |  |  |  |  | Inverted Part-Rearranged |  |  |  |  |
| --- | --- | --- | --- | --- | --- | --- | --- | --- | --- | --- | --- | --- | --- | --- | --- |
|  | Critical | % | Comp | % | Diff | Critical | % | Comp | % | Diff | Critical | % | Comp | % | Diff |
| Lamp | Lamp | 100.0 | Furniture | 18.8 | 81.3 | Keyhole | 46.9 | Vase | 9.4 | 37.5 | Vase | 28.1 | 3' / 'E' | 9.4 | 18.8 |
| Palm Tree | Palm tree | 100.0 | Monster | 12.5 | 87.5 | **Palm Tree / Tree** | **59.4** | Saw Blade | 15.6 | 43.8 | Cactus | 18.8 | Face | 15.6 | 3.1 |
| Rhino | Rhino | 100.0 | Ghost / Monster | 18.8 | 81.3 | Dinosaur | 18.8 | Dog | 18.8 | 0.0 | Person | 28.1 | Gargoyle | 21.9 | 6.3 |
| Elephant | Elephant | 96.9 | Landscape | 9.4 | 87.5 | **Elephant** | **90.6** | Person | 15.6 | 75.0 | **Elephant** | **50.0** | Mouth | 9.4 | 40.6 |
| Eagle | Eagle | 96.9 | Landscape | 9.4 | 87.5 | **Bird** | **18.8** | Face | 34.4 | -15.6 | Man with Hat | 37.5 | Person | 9.4 | 28.1 |
| Duck | Duck | 96.9 | Tree | 15.6 | 81.3 | **Duck** | **75.0** | Cliff | 9.4 | 65.6 | Person | 15.6 | Seahorse | 40.6 | -25.0 |
| Guitar | Guitar | 96.9 | Dock | 6.3 | 90.6 | Chess Piece | 15.6 | **Guitar** | **6.3** | 9.4 | Cloud | 9.4 | Gun | 6.3 | 3.1 |
| Hand | Hand | 96.9 | Waves | 9.4 | 87.5 | **Fingers / Hand** | **84.4** | Bird | 28.1 | 56.3 | **Fingers** | **56.3** | Claw | 15.6 | 40.6 |
| Train | Train | 96.9 | Faucet | 18.8 | 78.1 | Person | 50.0 | Gun | 25.0 | 25.0 | Faucet | 21.9 | Face | 25.0 | -3.1 |
| Mickey Mouse* | Mickey Mouse | 96.9 | Waves | 6.3 | 90.6 | **Mickey Mouse** | **34.4** | Landscape | 9.4 | 25.0 | Clown | 25.0 | Knife | 6.3 | 18.8 |
| Trumpet | Trumpet | 96.9 | **Instrument** | **15.6** | 81.3 | **Instrument** | **81.3** | Guitar | 21.9 | 59.4 | **Instrument** | **56.3** | **Instrument** | **37.5** | 18.8 |
| Boot | Boot | 93.8 | Face | 37.5 | 56.3 | **Shoe** | **56.3** | Mouth | 12.5 | 43.8 | Mouth | 12.5 | Lips | 9.4 | 3.1 |
| Flower | Flower | 93.8 | Person | 6.3 | 87.5 | **Flower** | **31.3** | Rhino | 6.3 | 25.0 | **Plant** | **50.0** | Leaf | 9.4 | 40.6 |
| Owl | Owl | 93.8 | Wave | 6.3 | 87.5 | **Bird** | **40.6** | Person | 12.5 | 28.1 | **Bird** | **50.0** | Monster | 18.8 | 31.3 |
| Pineapple | Pineapple | 93.8 | Wave | 6.3 | 87.5 | Clouds | 21.9 | Leaf | 12.5 | 9.4 | Berries | 18.8 | Leaf | 31.3 | -12.5 |
| Foot | Foot | 93.8 | Stalactites / Icicles | 15.6 | 78.1 | Baby | 34.4 | Scarf | 12.5 | 21.9 | Hair | 12.5 | Plant | 18.8 | -6.3 |
| Butterfly | Butterfly | 93.8 | Mountainside | 9.4 | 84.4 | **Butterfly / Wings** | **37.5** | Brass Instrument | 15.6 | 21.9 | **Butterfly** | **15.6** | Trumpet | 12.5 | 3.1 |
| House | House | 93.8 | Steam Whistle | 15.6 | 78.1 | Nose | 12.5 | Diving Board | 12.5 | 0.0 | Shelf | 15.6 | Heartbeat Signal | 6.3 | 9.4 |
| Face | Face | 90.6 | Vase | 15.6 | 75.0 | **Face** | **59.4** | Vase | 18.8 | 40.6 | **Face** | **25.0** | **Face** | **71.9** | -46.9 |
| Faucet | Faucet | 90.6 | Face | 18.8 | 71.9 | **Faucet** | **81.3** | Puzzle Piece | 12.5 | 68.8 | **Faucet** | **53.1** | Puzzle Piece | 12.5 | 40.6 |
| Snowman | Snowman | 90.6 | Waves | 6.3 | 84.4 | Bird | 28.1 | Bridge | 6.3 | 21.9 | Cloud | 28.1 | Waves | 6.3 | 21.9 |
| Toilet | Toilet | 90.6 | Mouth | 9.4 | 81.3 | Sink | 34.4 | Building | 9.4 | 25.0 | Shoe | 59.4 | Desk | 15.6 | 43.8 |
| Tree | Tree | 90.6 | Rock formation | 28.1 | 62.5 | Mountain | 21.9 | Mountain | 18.8 | 3.1 | Mountains | 21.9 | Mountain | 25.0 | -3.1 |
| Watering Can | Watering Can | 90.6 | Person | 9.4 | 81.3 | **Watering Can** | **50.0** | Tool | 9.4 | 40.6 | **Spout** | **59.4** | Mouth | 15.6 | 43.8 |
| Umbrella | Umbrella | 90.6 | Cat | 21.9 | 68.8 | **Umbrella** | **87.5** | Ocean | 12.5 | 75.0 | **Umbrella** | **68.8** | Mouth | 18.8 | 50.0 |
|  | Critical | % | Comp | % | Diff | Critical | % | Comp | % | Diff | Critical | % | Comp | % | Diff |
| Woman | Woman | 87.5 | Waves | 9.4 | 78.1 | Lamp |  | Plant | 12.5 | 30.5 | Vase | 15.6 | Person | 12.5 | 3.1 |
| Anchor | Anchor | 84.4 | Puzzle Piece | 12.5 | 71.9 | Tree | 28.1 | Mouth | 25.0 | 3.1 | Tree | 43.8 | Face | 12.5 | 31.3 |
| Axe | Axe | 84.4 | Hand/Fingers | 15.6 | 68.8 | Axe | 34.4 | Anvil | 15.6 | 18.8 | Axe | 53.1 | Corkscrew | 18.8 | 34.4 |
| Dog | Dog | 84.4 | Face | 12.5 | 71.9 | Mountain | 15.6 | Mountain | 25.0 | -9.4 | Person | 43.8 | Mountain | 25.0 | 18.8 |
| Seahorse | Seahorse | 84.4 | Tree | 12.5 | 71.9 | Seahorse | 25.0 | Winged Animal | 21.9 | 3.1 | Praying People | 28.1 | Dragon | 18.8 | 9.4 |
| Cow | Cow | 81.3 | Face | 12.5 | 68.8 | Mouth | 37.5 | Dog | 12.5 | 25.0 | Mouth | 18.8 | Mouth | 6.3 | 12.5 |
| Lightbulb | Lightbulb | 81.3 | Vase | 15.6 | 65.6 | Vase | 46.9 | Knife | 15.6 | 31.3 | Breasts | 12.5 | Vase | 18.8 | -6.3 |
| Bell | Bell | 78.1 | Vase / Urn | 18.8 | 59.4 | Lamp | 68.8 | Person | 15.6 | 53.1 | Lamp | 15.6 | Lamp | 25.0 | -9.4 |
| Fire Hydrant | Fire Hydrant | 78.1 | Traffic Light | 9.4 | 68.8 | Smokestack | 12.5 | Building | 31.3 | -18.8 | Key | 25.0 | Building | 28.1 | -3.1 |
| Teapot | Teapot | 75.0 | Bearded Man | 21.9 | 53.1 | Tree | 21.9 | Face | 6.3 | 15.6 | Person / Child | 53.1 | Mountain | 18.8 | 34.4 |
| Wine Glass | Wine Glass | 75.0 | Cleaver | 9.4 | 65.6 | Wine Glass | 37.5 | Wood | 12.5 | 25.0 | Top Hat | 78.1 | Mouth | 18.8 | 59.4 |
| Maple Leaf | Maple leaf | 71.9 | Face | 21.9 | 50.0 | Crystals | 9.4 | Mountain | 21.9 | -12.5 | Leaf | 9.4 | Cityscape | 15.6 | -6.3 |
| Pig | Pig | 71.9 | Canyon | 6.3 | 65.6 | Alien | 12.5 | Pig | 18.8 | -6.3 | Plant | 15.6 | Pig | 56.3 | -40.6 |
| Spray Bottle | Spray Bottle | 68.8 | Person | 15.6 | 53.1 | Water Fountain | 34.4 | Cartoon / Face | 34.4 | 0.0 | Faucet | 21.9 | Face | 34.4 | -12.5 |
| Grapes | Grapes | 65.6 | Stairs | 6.3 | 59.4 | Clouds | 56.3 | Tree / Leaf | 46.9 | 9.4 | Clouds | 75.0 | Leaf | 59.4 | 15.6 |
| Turtle | Turtle | 56.3 | Cave | 6.3 | 50.0 | Rabbit | 34.4 | Knife | 9.4 | 25.0 | Turtle | 40.6 | Seahorse | 9.4 | 31.3 |
| Wrench | Wrench | 53.1 | Face | 18.8 | 34.4 | Rhino | 9.4 | Tree | 9.4 | 0.0 | Tree | 43.8 | Face | 21.9 | 21.9 |
| Bottle | Bottle | 40.6 | Column | 15.6 | 25.0 | Stove/Furnace | 18.8 | Glass | 18.8 | 0.0 | Lamp post | 25.0 | Bottle | 28.1 | -3.1 |
| Bear | Bear | 37.5 | Mountains | 9.4 | 28.1 | Feet | 9.4 | Cityscape | 12.5 | -3.1 | Crowd | 9.4 | Mountainside | 9.4 | 0.0 |
| Rabbit | Lips | 34.4 | Face | 25.0 | 9.4 | Face | 18.8 | Waves | 6.3 | 12.5 | Person | 18.8 | Waves | 9.4 | 9.4 |
| Jet | Man w long face | 34.4 | Gun | 12.5 | 21.9 | Nose | 25.0 | Mouth | 12.5 | 12.5 | Airplane | 15.6 | Tree | 12.5 | 3.1 |
| Apple | Apple | 31.3 | Neck | 31.3 | 0.0 | Chin | 18.8 | Hand | 9.4 | 9.4 | Nose | 12.5 | Waves | 9.4 | 3.1 |
| Pear | Guitar | 78.1 | Waves | 6.3 | 71.9 | Woman | 31.3 | Waves | 18.8 | 12.5 | Female Body | 46.9 | Stringed Instrument | 18.8 | 28.1 |
| Mean (48) |  | 81.3 |  | 14.0 | 67.3 |  | 37.9 |  | 16.2 | 21.7 |  | 32.5 |  | 19.9 | 12.6 |
| Mean (44) |  | 84.7 |  | 13.6 | 71.1 |  | 39.3 |  | 16.6 | 22.6 |  | 33.3 |  | 20.6 | 12.7 |

### Procedure

#### Programs

All 24 programs were created outside of AMT as HTML files using Javascript/CSS/HTML and the JQuery Javascript library (version 1.11.3, https://jquery.com), and were then copied as source code into AMT. Stimuli (i.e., instructions, bipartite displays) were hosted on Imgur (https://imgur.com); their URLs were referenced by the programs. In each program, 24 bipartite stimuli were shown (8 in each type of display); half of the critical sides of each type were black, and half were white; half were on the left and half were on the right. Each source stimulus was only presented in one of the three types of displays (*Upright Intact, Upright Part-Rearranged*, and *Inverted Part-Rearranged*) in each program. Two different groups of programs were published, each presenting 24 of the 48 source stimuli. There were 12 programs within each group of programs. Black/white contrast and left/right location of the critical sides were balanced, and in each program one third of the stimuli were presented in each type of display (*Upright Intact, Upright Part-Rearranged*, and *Inverted Part-Rearranged*). Thus, across the 12 programs in each of the two groups, every stimulus was shown equally often in each of its three display types, and within display type, equally often with the critical sides in black/white and on the left/right.

Eligible participants could access only one program per group. Each program was viewed by 8 participants and participants never viewed the same stimulus more than once. In total, 32 participants provided up to three responses for each of the critical and complementary sides of each configuration of each source stimulus. Of the 174 participants, 156 completed one program and provided responses for 24 of the bipartite stimuli; 18 participants completed two programs (in different groups) and provided responses for all 48 stimuli (16 of each type, no overlap in source stimulus).

Participants had up to one hour to complete the experiment (see footnote 2). Participants had to click a button to advance through the programs which were segmented into pages. The first page was a consent form that was approved by the Human Subjects Protections Program at the University of Arizona. Participants could continue onto the rest of the program only after they indicated that they had read the consent form and agreed to participate in the experiment. The second page was an instruction page. The instructions showed a sample trial, and informed participants to use the three response boxes on the right and left sides of the screen to list up to three familiar objects resembled by the corresponding regions of the bipartite display. Participants were told they could type an ‘x’ in the top response box if they did not see any familiar objects on that side. Participants could not proceed to the next trial (the next page) without entering something in the top response boxes on the left and right sides. Figure 2 shows a sample trial.

**Figure 2.**
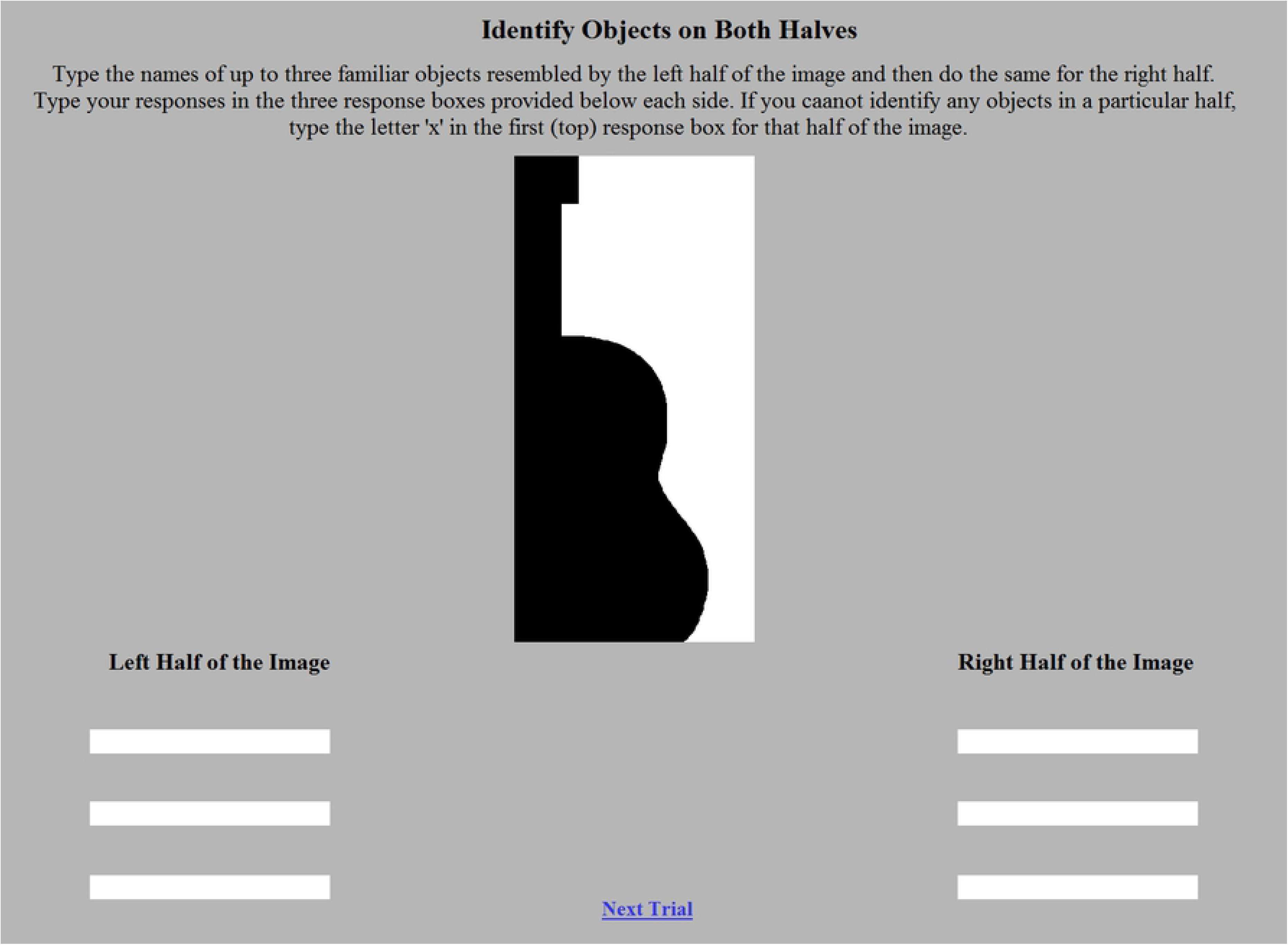
A sample trial from Experiment 1. Participants were presented with a bipartite stimulus; here, an *Upright Intact* version of the source stimulus “guitar” sketched in black on the left of the central border. Six response boxes were provided (three per side). They used these boxes to list any familiar objects resembled by each side of the stimulus. A button labelled ‘Next Trial’ would lead them to the next trial when they were ready.

After the instructions, participants completed 26 experimental trials: 24 trials with bipartite displays and two attention check trials. Of the 24 trials with bipartite displays, eight trials tested each of the three configuration types (upright Intact, upright Part-Rearranged, inverted Part-Rearranged). For each display type, the critical side was equally likely to be black or white, and located on the left or right within each program. On the two attention check trials, the bipartite stimulus was replaced with a white box. Inside the white box were written instructions on how to respond (e.g., “please write ‘fear’ in the top left and right box”). The attention check trials were included to make sure that participants were performing the task. If participants responded incorrectly on the attention check trials, their responses to the bipartite displays were discarded before they were viewed by an experimenter. The 26 experimental trials were presented in a random order. Time to complete each trial was unrestricted. After the experimental trials, participants were asked to provide any feedback or thoughts on a final page and were prompted to submit their responses.

## Data Analysis

Responses from all of the programs were sorted according to source stimulus and display type (*Upright Intact, Upright Part-Rearranged*, or *Inverted Part-Rearranged*), and bipartite stimulus side (critical or complementary). Responses to the critical and complementary sides were collapsed over contrast (black/white) and location relative to the central border (left/right). Responses were compiled across 32 participants (up to 96 responses per side given that participants could make up to three responses per side). Next, scorers cleaned up typing/spelling errors (e.g., consolidating ‘trumpet’ and ‘trumpit’) and grouped responses that seemed to denote similar object categories (e.g., ‘clarinet’ and ‘trumpet’ were grouped into single category response for the “Trumpet” source stimulus). These groupings were the basis for the inter-subject agreement scores (see below). Because participants differed in the level of specificity with which they identified objects resembled by the stimuli, responses made by different subjects were considered the same if they labeled the same basic-level object with a different name. For example, the responses ‘dwelling’ and ‘house’ made by different participants were both taken as evidence that the House source stimulus had been recognized at the basic level. If a single participant made two responses that were synonymous for a given region (i.e., ‘house’ and ‘dwelling’ as two different responses for the critical side of the border of the *Upright Intact* version of the House source stimulus), only one was counted. Each grouping of responses into one object category was initially made by a naïve scorer; their groupings were checked and confirmed by a second naïve scorer. Differences were referred to and resolved by the authors. A single object category could contain only one response per participant.

## Results

The best fitting object category perceived for a given side of the border of a given stimulus was selected as the one identified by the largest number of participants. Percent inter-subject agreement regarding this object category was determined by dividing the number of participants who made this response by 32 (the maximum number of responses if every participant contributed one response). Inter-subject agreement percentages are presented in Table 1. The identity of the source stimuli is listed in the left column. The three variants of the source stimuli – *Upright Intact*, *Upright Part-Rearranged*, and *Inverted Part Rearranged* – are arranged from left to right with five columns embedded under each type. These five columns list from left to right (1) the object category with the highest inter-subject agreement for the critical side of the central border, (2) the percent inter-subject agreement for that object category, (3) the object category with the highest inter-subject agreement for the complementary side of the central border, (4) the percent inter-subject agreement for that object category, and (5) the difference between the inter-subject agreement percentages for the critical and complementary sides of the border.

### Upright Intact displays

#### Critical side

Mean inter-subject agreement for the *Upright Intact* displays was 81.3%, indicating that on average the critical sides of the borders are good depictions of the source stimuli: The source stimuli are sorted by inter-subject agreement with one exception – the Pear stimulus is listed last because 78.1% of participants misidentified the critical side of the border as a “guitar.” The critical side of the upright Intact Jet display was also misidentified: 34.4% of participants identified it as a “face.” Given that inter-subject agreement was > 90% for the critical sides of source stimuli Guitar and Face, we recommend dropping these stimuli. We also recommend dropping the Rabbit stimulus because the critical side was identified as “lips” by the largest percentage of participants (34.4%) rather than as a rabbit.

#### Complementary side

The data indicate that the complementary sides of the borders of *Upright Intact* displays are not good depictions of well-known objects. Mean inter-subject agreement regarding objects denoted on the complementary side was 14.0%. We originally intended to use only bipartite displays in which participants indicated that the complementary side didn’t resemble anything familiar. In our early work, we found this was nearly impossible, so we instead set an upper cut-off of 23% inter-subject agreement on a single interpretation for portions of displays that could serve as complementary sides, because they depicted nominally “novel” objects [16]. We no longer set an a priori cutoff for the complementary sides: Inter-subject agreement for the object category resembled by the complementary side of the border of five of the 48 *Upright Intact* displays was ≥ 25% (source stimuli: Boot, Tree, Fire Hydrant, Rabbit, and Apple), although individual experimenters may choose to do so.

We note that the interpretations listed for the complementary side of the border of 14 of the *Upright Intact* displays were landscape features rather than objects: see responses of “landscape,” “waves,” “mountainside,” “rock formation,” building,” “canyon,” and “cave” (only “building” and “rock formation” generated ≥ 25% agreement). It is not clear whether past experience with landscape features influences figure assignment (although many of the landscape features named here do occlude other parts of a scene). Nevertheless, we list them in Table 1 because they produced the highest inter-subject agreement.

#### Critical – complementary difference

The difference between the inter-subject agreement for the critical and complementary sides of the border shown in the fifth column under *Upright Intact* displays was large – 67.3% on average. The *critical – complementary difference* was 0.00 for one of the source stimuli (Apple), however which leads us to suggest omitting this stimulus from the OMEFA II set.

#### Summary for Upright Intact displays

Based on the inter-subject agreement for the *Upright Intact* displays, we recommend omitting the last 4 stimuli on the list (Rabbit, Jet, Apple, and Pear). In what follows, we omit discussions of the data obtained for the other variants of these displays. That leaves a set of 44 *Upright Intact* displays, with mean inter-subject agreement of 84.7% for the critical side (range = 37.5% - 100%); 13.6% for the complementary side (range = 6.3% - 37.5%) and a mean *critical – complementary difference* of 71.1%, (range 25% - 90.6%)

### Upright Part-Rearranged Displays

#### Critical side

The mean inter-subject agreement regarding the category of the objects resembled by the critical side of the border of *Upright Part-Rearranged* displays was 39.3% (not counting the four source stimuli already rejected). This percentage indicates lower inter-subject agreement than for the *Upright Intact* displays, which we take as evidence that *Upright Part-Rearranged* displays are less likely to activate memory traces of well-known objects. For the critical sides of 18 of the *Upright Part-Rearranged* stimuli, however, the highest inter-subject agreement was for the same object category as identified for the critical sides of *Upright Intact* displays (see interpretations in bold). These responses are probably based on identification of a diagnostic part (e.g., the elephant’s trunk). The mean inter-subject agreement for these 18 stimuli (54.7%) was lower than for the corresponding *Upright Intact* stimuli (92.4%). The mean inter-subject agreement regarding the identity of the object denoted by the remaining 26 stimuli was low (28.6%) although not as low as for the complementary sides of these displays or of *Upright Intact* displays (see below). Perhaps configurations created from the parts of well-known objects resemble familiar objects more than configurations created from complements of those parts. For the critical side of the remaining 26 *Upright Part-Rearranged* stimuli, inter-subject agreement indicated that subjects perceived objects different from the source objects; for 11 of these displays inter-subject agreement was ≥ 25% (source stimuli: lamp, train, foot, snowman, toilet, woman, cow, light bulb, bell, spray bottle, grapes, turtle).

#### Complementary side

Mean inter-subject agreement regarding the category of the objects resembled by the complementary side of the border was low (16.6%), and approximately the same as for the complementary side of the *Upright Intact* displays. Inter-subject agreement was ≥ 25% for the complementary sides of eight of the source stimuli (Eagle, Hand, Train, Anchor, Dog, Fire Hydrant, Spray Bottle, and Grapes). Two of these interpretations (“mountain” and “building”) were landscape features rather than objects per se; one was the source object (Pig).

#### Critical – complementary differences

The mean difference between the inter-subject agreement for the critical and complementary sides of the border observed for *Upright Part-Rearranged* displays was 22.6%, quite a bit smaller than for *Upright Intact* displays. The c*ritical – complementary differences* were negative for five stimuli. Most of these negative differences were small; the largest negative difference (−18.8) was obtained when the inter-subject agreement for the complementary side was a landscape feature (“building”).

#### Summary for upright part-rearranged displays

For the current set of 44 stimuli, the mean inter-subject agreement was 39.4% for the critical side (range = 9.4% - 90.6%); 16.6% for the complementary side (range = 6.3% - 34.4%) and a mean *critical – complementary difference* of 22.6%, (range −18.8% - 75.0%). As manifested by participants’ explicit responses, overall, the critical sides of the *Upright Part-Rearranged* displays are less likely to activate traces of previously seen objects. Inter-subject agreement was higher for some of the displays, perhaps because of the presence of diagnostic parts.

### Inverted Part-Rearranged Displays

#### Critical side

The mean inter-subject agreement regarding well known objects resembled by the critical side of the border of *Inverted Part-Rearranged* displays (33.3%) was slightly lower than for the *Upright Part-Rearranged* displays and substantially lower than for the *Upright Intact* displays. We take these data as evidence that the critical sides of these stimuli do not highly activate memory traces of well-known objects. For 13 of the 44 *Inverted Part-Rearranged* stimuli, however, the largest percentage of participants identified the critical side as the source object. Once again, we hypothesize that this high inter-subject agreement is based on the identification of diagnostic parts: 11 of these 13 displays are a subset of the 17 *Upright Part-Rearranged* displays for which participants agreed in naming the source object on the critical side of the border. The mean inter-subject agreement for these 11 *Inverted Part-Rearranged* displays (51.1%) was substantially lower than for the corresponding *Upright Intact* displays (89.2%) but only slightly lower than for the corresponding *Upright Part-Rearranged* displays (61.4%). The mean inter-subject agreement for the critical sides of the remaining 31 *Inverted Part-Rearranged* displays was 28.3%.

#### Complementary side

Inter-subject agreement regarding the object category resembled by the complementary side of the border of the 44 stimuli under consideration was 20.6%. For five of the displays, the highest inter-subject agreement was for the source object (source objects: Face, Woman, Trumpet, Bottle, and Pig). For the Face and the Pig source stimuli, the inter-subject agreement regarding the source object interpretation for the complementary side of the border was higher than for the critical side of the border, leading to negative *critical – complementary* differences (see below). For nine other displays, inter-subject agreement that the complementary side of the border denoted a different object was ≥ 25% (Duck, Train, Pineapple, Tree, Dog, Bell, Fire Hydrant, Spray Bottle, and Grapes). Three of these interpretations were landscape features rather than objects, and two were simply parts (e.g., “leaf”).

##### Critical – complementary difference

The mean difference between the inter-subject agreement for the critical and complementary sides of the border observed for *Inverted Part-Rearranged* displays was 12.7%, smaller than for the *Upright Part-Rearranged* displays. The smaller difference was obtained because between-subjects agreement was both lower for the critical side and higher for the complementary side. The *critical – complementary differences* were negative for 13 stimuli. The largest negative difference (−46.9%) was obtained when the higher inter-subject agreement for the complementary side was for “face,” which may be a manifestation of pareidolia.

## Discussion

We present OMEFA-II, a new, high-resolution set of bipartite stimuli (N = 44 source stimuli) with normative data regarding explicit judgments of the familiar objects denoted on both the critical and complementary sides of the central borders of three different display types: *Upright Intact* displays (N = 44; 88 sides), *Upright Part-Rearranged* displays (N = 44; 88 sides), and *Inverted Part-Rearranged* displays (N = 44; 88 sides). These comprehensive norms allow the difference in inter-subject agreement for the critical and complementary sides to be calculated for 132 displays. The OMEFA-II displays and the normative data reported here will be valuable for experiments conducted with participants with brain damage as well as those with intact brains for investigating questions concerning parts and wholes, high-level influences on perception, and for tests of competitive models of perception.

The inter-subject agreement measured here is one means of quantifying the extent to which traces of previously seen objects are activated by different sides of a border, but this activation occurs implicitly during perceptual organization. Behavioral measures such as the probability of perceiving the figure on the critical side of the border, event-related potentials (ERPs), and the blood oxygen dependent (BOLD) response in fMRI experiments, perhaps in combination with multi-voxel pattern analysis (MVPA), may also quantify activation of traces of previously seen objects. Correlating the data presented in Table 1 with these indices may be fruitful in understanding object perception in general, figure assignment in particular, and any underlying competition between objects that might be perceived on opposite sides of a border.

In addition to inter-subject agreement for the critical and complementary sides of the border individually, we report the difference in inter-subject agreement regarding the objects sketched on the critical versus the complementary sides of the border. On current inhibitory competition accounts of figure assignment (e.g., [24, 25]), this difference may better predict whether a figure will be perceived on the critical side of a border than the inter-subject agreement regarding the critical side alone. The comprehensive set of norms presented here allows future experiments to test which is the better predictor.

Although inter-subject agreement is informative about *which* traces of previously seen objects are activated, they cannot assess *how* quickly they are activated. In previous research, substantially larger effects of familiar configuration were found for *Upright* than *Inverted Intact* displays (e.g., [2, 13, 16, 17]). This orientation-dependent difference has been attributed to the time required for evidence to accumulate in neural populations coding for the familiar object (longer for inverted than upright displays; [26]). The orientation-dependency of the familiar configuration prior has been taken to indicate that priors for figure assignment must be available quickly in order to influence figure assignment (for review see [15]). Indeed, once the critical sides of *Inverted Intact* displays are perceived as figures, the familiar objects they portray can often be identified. (This is why we did not obtain norms for the critical side of *Inverted Intact* displays.) Nevertheless, knowing that critical sides depict inverted familiar objects does not increase the likelihood of seeing the figure on the side where an inverted version of the intact object is sketched [13].

For some of the *Upright* and *Inverted Part-Rearranged* displays, sizeable inter-subject agreement seemed to be based on diagnostic parts. In future research it will be interesting to test whether access to object categories via diagnostic parts as evidenced by these explicit responses generated while the stimuli were exposed for long durations is sufficient for past experience effects on figure assignment. Given the large set normed here this can be done for *Upright Part-Rearranged* displays by comparing performance with the 18 displays for which the largest percentage of participants identified the source stimulus and the remaining 26 displays for which the largest percentage of participants did not identify the source stimulus. (For the *Inverted Part-Rearranged* displays, this would be a comparison between the 13 and 31 displays for which the largest percentage of participants did and did not identify the source stimulus). Previous studies have shown that the critical side of the border is substantially and significantly less likely to be perceived as the figure in *Upright Part-Rearranged* displays than *Upright Intact* displays (e.g., [13, 16, 18–20]). Yet none of those experiments used the large set of stimuli normed here that affords a sensitive analysis of differences within the set of *Upright Part-Rearranged* displays based on whether diagnostic parts supported identification of the source stimulus. (We note that 13 of the 18 stimuli for which inter-subject agreement was highest that the critical side of the border resembled the source object category are new stimuli that were not previously normed.)

Some of the interpretations that garnered >25% agreement were landscape features rather than objects. A small percentage of similar responses was observed in previous norming studies, but they did not exceed 25% agreement. It could be interesting to test whether, for an equivalent level of inter-subject agreement, landscape features and concrete objects are equivalent priors for figure assignment.

## Conclusion

We present normative data obtained for an expanded set of bipartite stimuli that are well-suited for assessing high-level influences on figure assignment, an essential component of object perception – the OMEFA-II stimulus set. The bipartite stimuli are divided into two equal area regions by a central border. Normative data were obtained by presenting the bipartite stimuli to AMT participants who were asked to identify any familiar objects sketched by central border on both a critical side and a complementary side. The critical side depicted either an intact version of an upright familiar object (*Upright Intact* displays), or a part-rearranged version in an upright or inverted orientation (*Upright Part-Rearranged* and *Inverted Part-Rearranged* displays respectively). The stimuli, as well as Excel files of Table 1, Appendix A, the AMT data sorted by stimulus type (and within stimulus type by critical and complementary side), and the full data set are available online (https://osf.io/j9kz2/).

---

**Appendix A.**
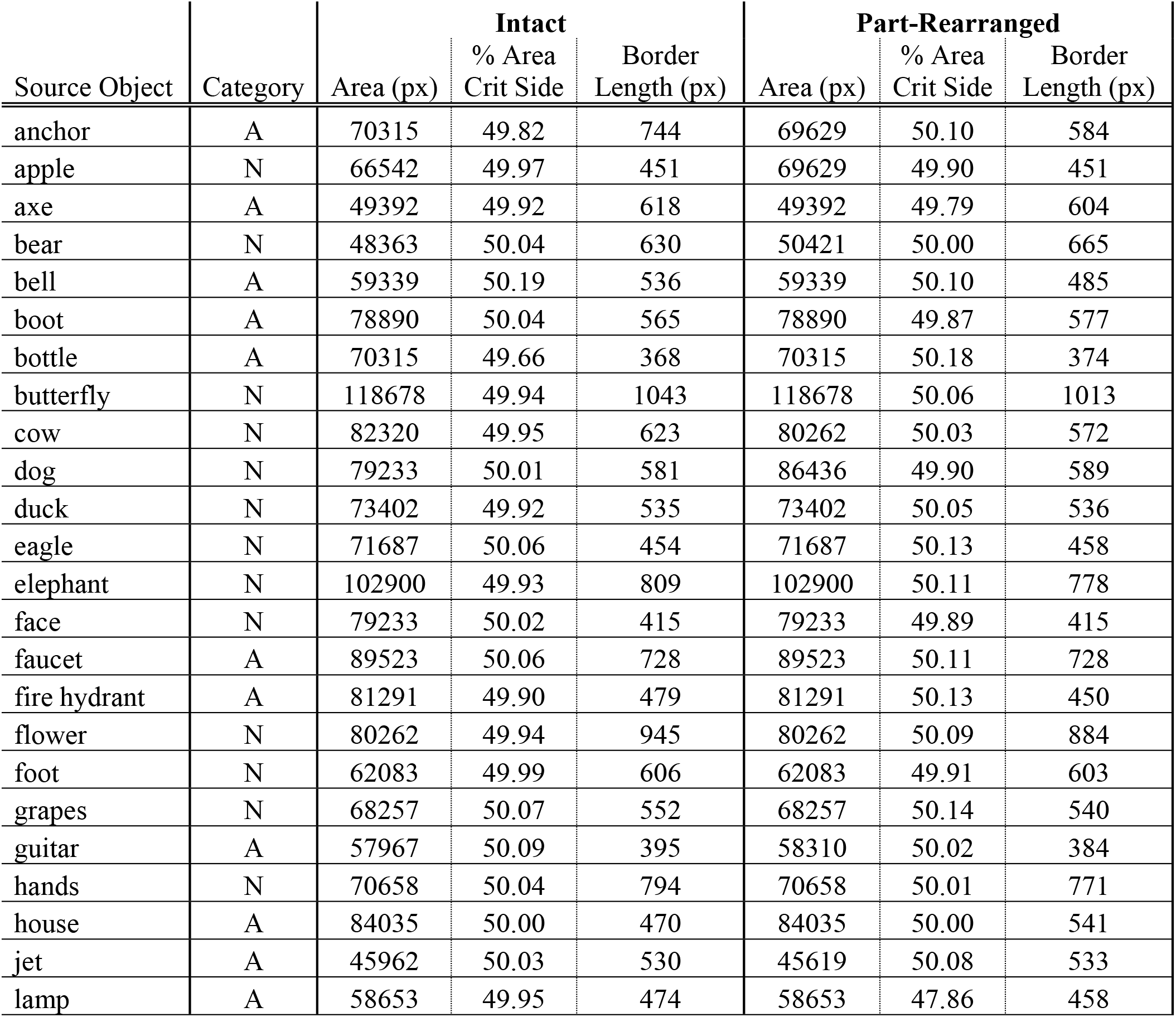
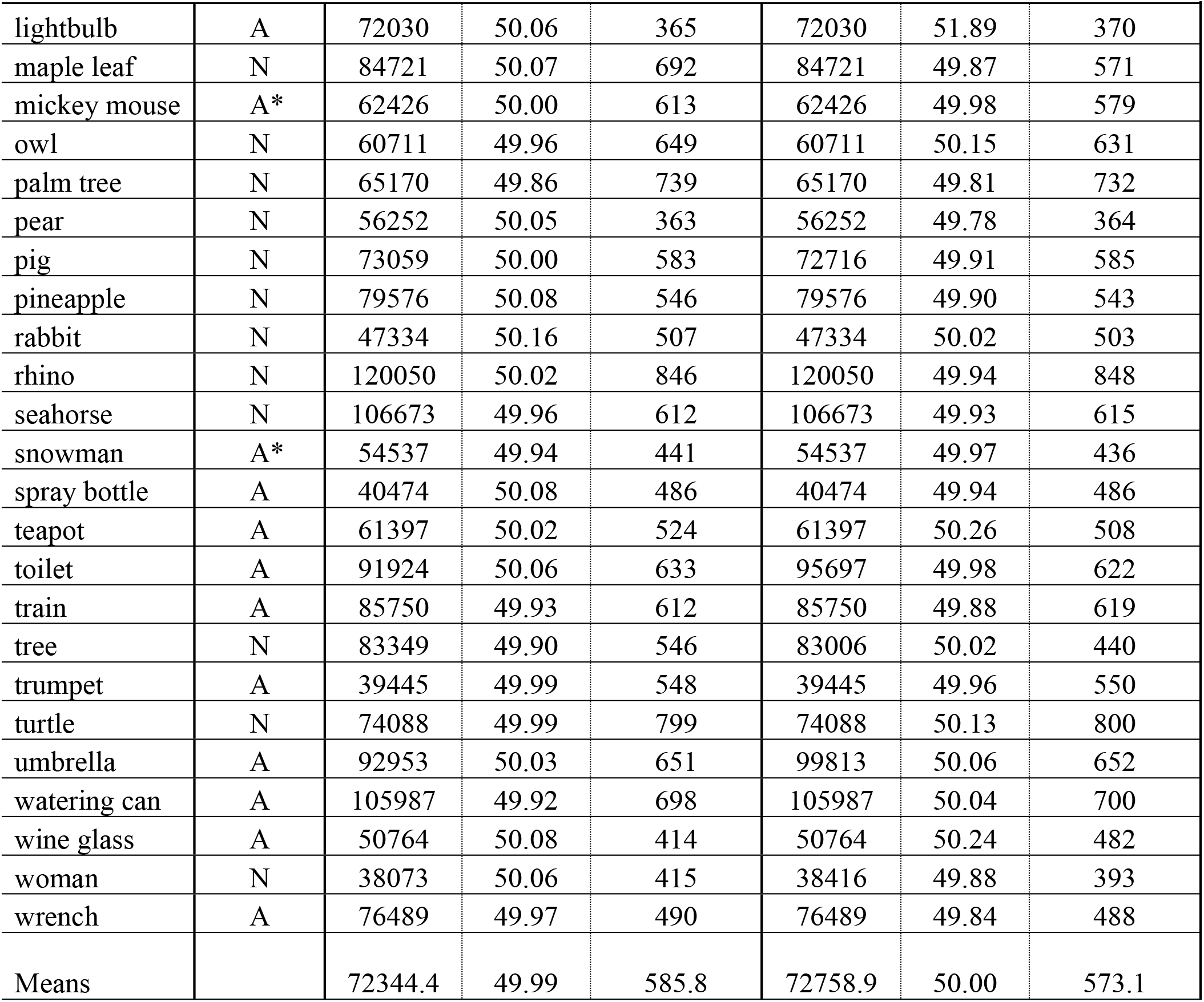
Image statistics for bipartite images. Statistics are shown for *Intact* and *Part-Rearranged* displays. The statistics are the same for both *Upright* and *Inverted* orientations. Category denotes whether the source object is Natural (N) or Artificial (A); * = ambiguous. “Area (px)” is the total number of pixels in the display. “% Area Crit Side” is the percentage of pixels on the critical side of the border. The percentage of pixels on the complementary side of the border is (1 – “% Area Crit Side.” “Border Length” is the length of the central border in pixels calculated using the bwperim function in MATLAB (2016b; MathWorks, Natick, MA).

| Source Object | Category | Intact |  |  | Part-Rearranged |  |  |
| --- | --- | --- | --- | --- | --- | --- | --- |
|  |  | Area (px) | % Area Crit Side | Border Length (px) | Area (px) | % Area Crit Side | Border Length (px) |
| anchor | A | 70315 | 49.82 | 744 | 69629 | 50.10 | 584 |
| apple | N | 66542 | 49.97 | 451 | 69629 | 49.90 | 451 |
| axe | A | 49392 | 49.92 | 618 | 49392 | 49.79 | 604 |
| bear | N | 48363 | 50.04 | 630 | 50421 | 50.00 | 665 |
| bell | A | 59339 | 50.19 | 536 | 59339 | 50.10 | 485 |
| boot | A | 78890 | 50.04 | 565 | 78890 | 49.87 | 577 |
| bottle | A | 70315 | 49.66 | 368 | 70315 | 50.18 | 374 |
| butterfly | N | 118678 | 49.94 | 1043 | 118678 | 50.06 | 1013 |
| cow | N | 82320 | 49.95 | 623 | 80262 | 50.03 | 572 |
| dog | N | 79233 | 50.01 | 581 | 86436 | 49.90 | 589 |
| duck | N | 73402 | 49.92 | 535 | 73402 | 50.05 | 536 |
| eagle | N | 71687 | 50.06 | 454 | 71687 | 50.13 | 458 |
| elephant | N | 102900 | 49.93 | 809 | 102900 | 50.11 | 778 |
| face | N | 79233 | 50.02 | 415 | 79233 | 49.89 | 415 |
| faucet | A | 89523 | 50.06 | 728 | 89523 | 50.11 | 728 |
| fire hydrant | A | 81291 | 49.90 | 479 | 81291 | 50.13 | 450 |
| flower | N | 80262 | 49.94 | 945 | 80262 | 50.09 | 884 |
| foot | N | 62083 | 49.99 | 606 | 62083 | 49.91 | 603 |
| grapes | N | 68257 | 50.07 | 552 | 68257 | 50.14 | 540 |
| guitar | A | 57967 | 50.09 | 395 | 58310 | 50.02 | 384 |
| hands | N | 70658 | 50.04 | 794 | 70658 | 50.01 | 771 |
| house | A | 84035 | 50.00 | 470 | 84035 | 50.00 | 541 |
| jet | A | 45962 | 50.03 | 530 | 45619 | 50.08 | 533 |
| lamp | A | 58653 | 49.95 | 474 | 58653 | 47.86 | 458 |
|  |  | Area (px) | % Area<br>Crit Side | Border<br>Length (px) | Area (px) | % Area<br>Crit Side | Border<br>Length (px) |
| lightbulb | A | 72030 | 50.06 | 365 | 72030 | 51.89 | 370 |
| maple leaf | N | 84721 | 50.07 | 692 | 84721 | 49.87 | 571 |
| mickey mouse | A* | 62426 | 50.00 | 613 | 62426 | 49.98 | 579 |
| owl | N | 60711 | 49.96 | 649 | 60711 | 50.15 | 631 |
| palm tree | N | 65170 | 49.86 | 739 | 65170 | 49.81 | 732 |
| pear | N | 56252 | 50.05 | 363 | 56252 | 49.78 | 364 |
| pig | N | 73059 | 50.00 | 583 | 72716 | 49.91 | 585 |
| pineapple | N | 79576 | 50.08 | 546 | 79576 | 49.90 | 543 |
| rabbit | N | 47334 | 50.16 | 507 | 47334 | 50.02 | 503 |
| rhino | N | 120050 | 50.02 | 846 | 120050 | 49.94 | 848 |
| seahorse | N | 106673 | 49.96 | 612 | 106673 | 49.93 | 615 |
| snowman | A* | 54537 | 49.94 | 441 | 54537 | 49.97 | 436 |
| spray bottle | A | 40474 | 50.08 | 486 | 40474 | 49.94 | 486 |
| teapot | A | 61397 | 50.02 | 524 | 61397 | 50.26 | 508 |
| toilet | A | 91924 | 50.06 | 633 | 95697 | 49.98 | 622 |
| train | A | 85750 | 49.93 | 612 | 85750 | 49.88 | 619 |
| tree | N | 83349 | 49.90 | 546 | 83006 | 50.02 | 440 |
| trumpet | A | 39445 | 49.99 | 548 | 39445 | 49.96 | 550 |
| turtle | N | 74088 | 49.99 | 799 | 74088 | 50.13 | 800 |
| umbrella | A | 92953 | 50.03 | 651 | 99813 | 50.06 | 652 |
| watering can | A | 105987 | 49.92 | 698 | 105987 | 50.04 | 700 |
| wine glass | A | 50764 | 50.08 | 414 | 50764 | 50.24 | 482 |
| woman | N | 38073 | 50.06 | 415 | 38416 | 49.88 | 393 |
| wrench | A | 76489 | 49.97 | 490 | 76489 | 49.84 | 488 |
| Means |  | 72344.4 | 49.99 | 585.8 | 72758.9 | 50.00 | 573.1 |

## References

[1] Hochberg J. Perception I: Color and shape. In: Kling JW, Riggs LA, editors. Woodworth and Schlossberg’s experimental psychology. New York, NY; Holt, Rinehart and Winston, 1971:395–474.

[2] Peterson MA, Gibson BS. Must figure-ground organization precede object recognition? An assumption in peril. Psychol Sci. 1994;5:253–9. doi: 10.1111/j.1467-9280.1994.tb00622.x

[3] Pomerantz JR, Kubovy M. Theoretical approaches to perceptual organization: Simplicity and likelihood principles. In: Boff KR, Kaufman L, Thomas JP, editors. Handbook of perception and human performance. New York, NY: Wiley; 1986;36:36–1-46.

[4] Rubin, E. Figure and ground. In Beardslee DC, Wertheimer M, editors. Readings in perception. Princeton, NJ: Van Nostrand, 1915/1958:194–203.

[5] Hulleman J, Humphreys GW. A new cue to figure-ground coding: Top-bottom polarity. Vision Res. 2004;44:2779–91. doi: 10.1016/j.visres.2004.06.012

[6] Kanizsa G, Gerbino W. Convexity and Symmetry in figure-ground organization. In Henle M, editor. Vision and artifact. New York, NY: Springer, 1976:25–33.

[7] Mojica AJ, Peterson MA. Display-wide influences on figure-ground perception: The case of symmetry. Atten Percept Psychophys. 2014;76:1069–84. doi: 10.3758/s13414-014-0646-y

[8] O’Shea RP, Blackburn SG, Ono H. Contrast as a depth cue. Vision Res. 1994;34:1595–604. doi: 10.1016/0042-6989(94)90116-3

[9] Peterson MA, Salvagio E. Inhibitory competition in figure-ground perception: Context and convexity. J Vis. 2008;8:1–13. doi: 10.1167/8.16.4

[10] Vecera SP, Vogel EK, Woodman GF. Lower region: A new cue for figure-ground assignment. J Exp Psycho Gen. 2002;131:194–205. doi: 10.1037/0096-3445.131.2.194

[11] Peterson MA, Kimchi R. Perceptual organization in vision. In: Reisberg D, editor. The Oxford handbook of cognitive psychology. Oxford University Press, 2013:9–31.

[12] Peterson MA, Gibson BS. The initial identification of figure-ground relationships: Contributions from shape recognition processes. Bull Psychon Soc. 1991;29:199–202. doi: 10.3758/BF03335234

[13] Peterson MA, Harvey EM, Weidenbacher HJ. Shape recognition contributions to figure-ground reversal: Which route counts?. J Exp Psychol Hum Percept Perform. 1991;17:1075–89. doi: 10.1037/0096-1523.17.4.1075

[14] Peterson MA. Object recognition processes can and do operate before figure–ground organization. Curr Dir Psychol Sci. 1994;3:105–11. doi: 10.1111/1467-8721.ep10770552

[15] Peterson MA. Past experience and meaning affect object detection: A hierarchical Bayesian approach. In Federmeier KD, Beck DM, editors. Psychology of learning and motivation: Knowledge and vision. Cambridge, MA: Academic Press, 2019;70:223.

[16] Gibson BS, Peterson MA. Does orientation-independent object recognition precede orientation-dependent recognition? Evidence from a cuing paradigm. J Exp Psychol Hum Percept Perform. 1994;20:299. doi: 10.1037/0096-1523.20.2.299

[17] Peterson MA, Gibson BS. Object recognition contributions to figure-ground organization: Operations on outlines and subjective contours. Percept Psychophys. 1994;56:551–64. doi: 10.1111/j.1467-9280.1994.tb00622.x

[18] Barense MD, Ngo JK, Hung LH, Peterson MA. Interactions of memory and perception in amnesia: The figure–ground perspective. Cereb Cortex. 2012;22:2680–91. doi: 10.1093/cercor/bhr347

[19] Peterson MA, De Gelder B, Rapcsak SZ, Gerhardstein PC, Bachoud-Lévi AC. Object memory effects on figure assignment: Conscious object recognition is not necessary or sufficient. Vision Res. 2000;40:1549–67. doi: 10.1016/S0042-6989(00)00053-5

[20] Peterson MA, Gerhardstein PC, Mennemeier M, Rapcsak SZ. Object-centered attentional biases and object recognition contributions to scene segmentation in left-and right-hemisphere-damaged patients. Psychobiology. 1998;26:357–70. doi: 10.3758/BF03330622

[21] Cacciamani L, Wager E, Peterson MA, Scalf PE. Age-related changes in perirhinal cortex sensitivity to configuration and part familiarity and connectivity to visual cortex. Front Aging Neurosci. 2017;9:291. doi: 10.3389/fnagi.2017.00291

[22] Peterson MA, Cacciamani L, Mojica AJ, Sanguinetti JL. Meaning can be accessed for the ground side of a figure. Gestalt Theory. 2012;34:297–314.

[23] Peer E, Vosgerau J, Acquisti A. Reputation as a sufficient condition for data quality on Amazon Mechanical Turk. Behav Res Methods. 2014;46:1023–31. doi: 10.3758/s13428-013-0434-y

[24] Kogo N, Strecha C, Van Gool L, Wagemans J. Surface construction by a 2-D differentiation–integration process: A neurocomputational model for perceived border ownership, depth, and lightness in Kanizsa figures. Psychol Rev. 2010;117:406–39. doi: 10.1037/a0019076

[25] Vecera SP, O’reilly RC. Figure-ground organization and object recognition processes: an interactive account. J Exp Psychol Hum Percept Perform. 1998;24:441–62. doi: 10.1037/0096-1523.24.2.441

[26] Perrett DI, Oram MW, Ashbridge E. Evidence accumulation in cell populations responsive to faces: an account of generalisation of recognition without mental transformations. Cognition. 1998;67:111–45. doi: 10.1016/S0010-0277(98)00015-8

